# Structure-based virtual screening identifies small molecule inhibitors of O-fucosyltransferase SPINDLY

**DOI:** 10.1101/2023.06.13.544843

**Authors:** Yalikunjiang Aizezi, Hongming Zhao, Zhenzhen Zhang, Yang Bi, Qiuhua Yang, Guangshuo Guo, Hongliang Zhang, Hongwei Guo, Kai Jiang, Zhi-Yong Wang

## Abstract

Protein O-glycosylation is a nutrient-signaling mechanism that plays essential roles in maintaining cellular homeostasis across different species. In plants, SPINDLY (SPY) and SECRET AGENT (SEC) catalyze posttranslational modifications of hundreds of intracellular proteins by O-fucose and O-linked N-acetylglucosamine, respectively. SPY and SEC play overlapping roles in cellular regulation and loss of both SPY and SEC causes embryo lethality in Arabidopsis. Using structure-based virtual screening of chemical libraries followed by *in vitro* and *in planta* assays, we identified a <u>S</u>PY <u>O</u>-<u>f</u>ucosyltransferase <u>i</u>nhibitor (SOFTI). Computational analyses predicted that SOFTI binds to the GDP-fucose-binding pocket of SPY and competitively inhibits GDP-fucose binding. *In vitro* assays confirmed that SOFTI interacts with SPY and inhibits its O-fucosyltransferase activity. Docking analysis identified additional SOFTI analogs that showed stronger inhibitory activities. SOFTI treatment of Arabidopsis seedlings decreased protein O-fucosylation and caused phenotypes similar to the *spy* mutants, including early seed germination, increased root hair density, and defect in sugar-dependent growth. By contrast, SOFTI had no visible effect on the *spy* mutant. Similarly, SOFTI inhibited sugar-dependent growth of tomato seedlings. These results demonstrate that SOFTI is a specific SPY O-fucosyltransferase inhibitor and a useful chemical tool for functional studies of O-fucosylation and potentially for agricultural management.

## Introduction

Like phosphorylation, O-glycosylation of nuclear and cytoplasmic proteins plays a crucial role in cellular signaling and regulation. O-linked N-acetylglucosamine (O-GlcNAc) modification catalyzed by O-GlcNAc transferase (OGT) is known to mediate nutrient sensing and regulate a variety of biological processes in animals (Hart et al., 2007). While there is only one OGT in mammals, the Arabidopsis genome encodes two OGT homologs, SPINDLY (SPY) and SECRET AGENT (SEC). Genetic studies indicated overlapping functions of SPY and SEC in a wide range of developmental processes including shared functions essential for viability. SPY and SEC catalyze protein O-fucosylation and O-GlcNAcylation, respectively, utilizing GDP-fucose and UDP-GlcNAc as their respective donor substrates (Zentella et al., 2016; Zentella et al., 2017; Sun, 2021). Hundreds of nuclear and cytoplasmic proteins with regulatory roles are modified by O-fucose and O-GlcNAc; how the modifications regulate their functions remains largely unknown (Xu et al., 2017; Bi et al., 2023; Zentella et al., 2023).

The Arabidopsis *spindly* (*spy*) mutant was identified for its gibberellin (GA) hypermorphic phenotypes, including slender shoot organs, GA-independent seed germination, and early flowering. Subsequent studies uncovered *spy’s* GA-independent phenotypes including cytokinin hyporesponse, aberrant trichome branching and root hair patterning, altered inflorescence phyllotaxis, reduced fertility (Silverstone et al., 2007), and sugar-promoted seedling growth (Bi et al., 2023). In contrast, loss of SEC results in rather mild phenotypes, showing decreased sensitivity to plant hormone GA and early flowering under short-day conditions (Zentella et al., 2016; Xing et al., 2018). The *spy sec* double mutant is embryonic lethal, which indicates their importance in an essential process and makes it difficult to dissect their shared functions (Hartweck et al., 2002).

SPY/O-fucosylation and SEC/O-GlcNAcylation regulate the functions of several key proteins. One of the most thoroughly studied examples is DELLA, a growth repressor protein that negatively regulates GA signaling. DELLA is antagonistically regulated by O-fucose and O-GlcNAc modifications, which respectively enhance and inhibit its interaction with downstream transcription factors, including BZR1, PIF3, and PIF4 (Zentella et al., 2016; Zentella et al., 2017). SPY facilitates cytokinin signaling by stabilizing the TCP14 and TCP15 transcription factors (Steiner et al., 2012; Steiner et al., 2016). SPY also plays a role in the regulation of alternative RNA splicing events by O-fucosylation of ACINUS, a component of the splicing complex that is involved in the regulation of seed germination, flowering, and ABA signal transduction (Bi et al., 2021). Furthermore, SPY O-fucosylates PRR5 to regulate circadian rhythms (Wang et al., 2020).

Consistent with the redundant functions of SPY and SEC for viability, proteomic studies revealed significant overlaps between proteins modified by O-GlcNAc or O-fucose (Bi et al., 2023; Zentella et al., 2023). For example, about 49% (128/262) of the O-GlcNAcylated proteins are modified by O-fucose (Bi et al., 2023). Many O-GlcNAcylated proteins are also substrates of the BIN2 kinase, a key component of the brassinosteroid signaling pathway, whereas the O-fucosylated proteins tend to be targets of the Target of Rapamycin (TOR) kinase, which mediate nutrient signaling (Kim et al., 2023; Bi et al., 2023). Thus, the proteomic data indicates that the nutrient-and hormone-signaling pathways crosstalk through O-glycosylation and phosphorylation of common target proteins. Considering the lethality of the *spy sec* double mutant, elucidating the independent and shared functions of SPY or SEC requires conditional inactivation of each protein.

In animals, small molecule inhibitors of OGT have been instrumental in the functional study of O-GlcNAc modification (Alteen et al., 2021). Small molecules have also advanced many aspects of plant biology (Lepri et al., 2023). However, chemical inhibitors are yet to be identified for functional study of O-glycosylation in plants. Traditionally, identification of small molecule inhibitors involves phenotype- or assay-based screening of large chemical libraries (Lepri et al., 2023). Structure-based virtual screening has been widely used in human drug discovery research but requires three-dimensional structure of the target protein and has not been widely used in plant research. Recently, a breakthrough in computational structure prediction has generated structures of all proteins of Arabidopsis and major crops (Varadi et al., 2022). Whether the predicted structures are reliable for virtual screening for chemical binders remains arguable (Wong et al., 2022; Scardino et al., 2023).

Here we identify small molecule inhibitors of SPY through structure-based virtual screening based on the SPY structure predicted by AlphaFold (Jumper et al., 2021). Following computational docking of over 13,890 compounds, we tested the top candidates by *in vitro* binding and enzyme inhibition assays as well as *in vivo* treatment of plants. We identified small molecule compounds as SPY O-fucosyltransferase inhibitors (SOFTIs). We show that SOFTI treatment reduces protein O-fucosylation in Arabidopsis seedlings and causes phenotypes that resemble the *spy* mutants. Our study demonstrates the promise of AlphaFold-enabled virtual screening and identifies powerful tools for the investigation of protein O-fucosylation.

## Results

### Combination of structure-based virtual screening and *in vitro* binding assays identified 6 SPY binders

To find small molecule inhibitors of SPY, we performed structure-based virtual screening for compounds that fit the ligand binding pocket of SPY using the AlphaFold-predicted structure of SPY (uniport ID: A0A654F7U9). A library containing 13,890 compounds with broad diversity in chemical scaffolds or bioactivities was screened using Molecular Operating Environment (MOE), a commercial molecular docking software. MOE calculates the affinity of a protein-ligand interaction into a numerical value called S-score. The compounds were further selected according to the following criteria: 1. an S-score lower than -6; 2. predicted to form more than three polar interactions with the amino acid residues within the SPY ligand-binding pocket; 3. adopt a relatively simple structure for better cell permeability and convenience for future structural optimization. The virtual screening identified 130 compounds that meet the criteria and thus were considered candidates for inhibitors that compete with the donor substrate GDP-fucose.

To narrow down these candidate compounds, we tested the binding of 130 compounds to the full-length recombinant SPY protein produced from *E. coli* (Supplementary Figure 1) using the protein thermal shift assay (TSA). Six of these compounds (SPI-1 to SPI-6) affected the thermal stability of SPY by more than 3 degrees at 50 μM concentration (Figure 1B), indicating their direct binding to SPY. We further tested their binding to SPY using Bio-layer interferometry (BLI) (Abdiche et al., 2008), which measures the binding kinetics between SPY and the compounds. One of these components showed a concentration-dependent effect on the melting temperature (T_m_) of SPY (Figure 1C-E) and a dissociation constant (K_d_) value of 37 μM in BLI assay (Figure 1F). We named this compound SOFTI (<u>S</u>PINDLY <u>O</u>-fucosyltransferase inhibitor).

**Figure 1.**
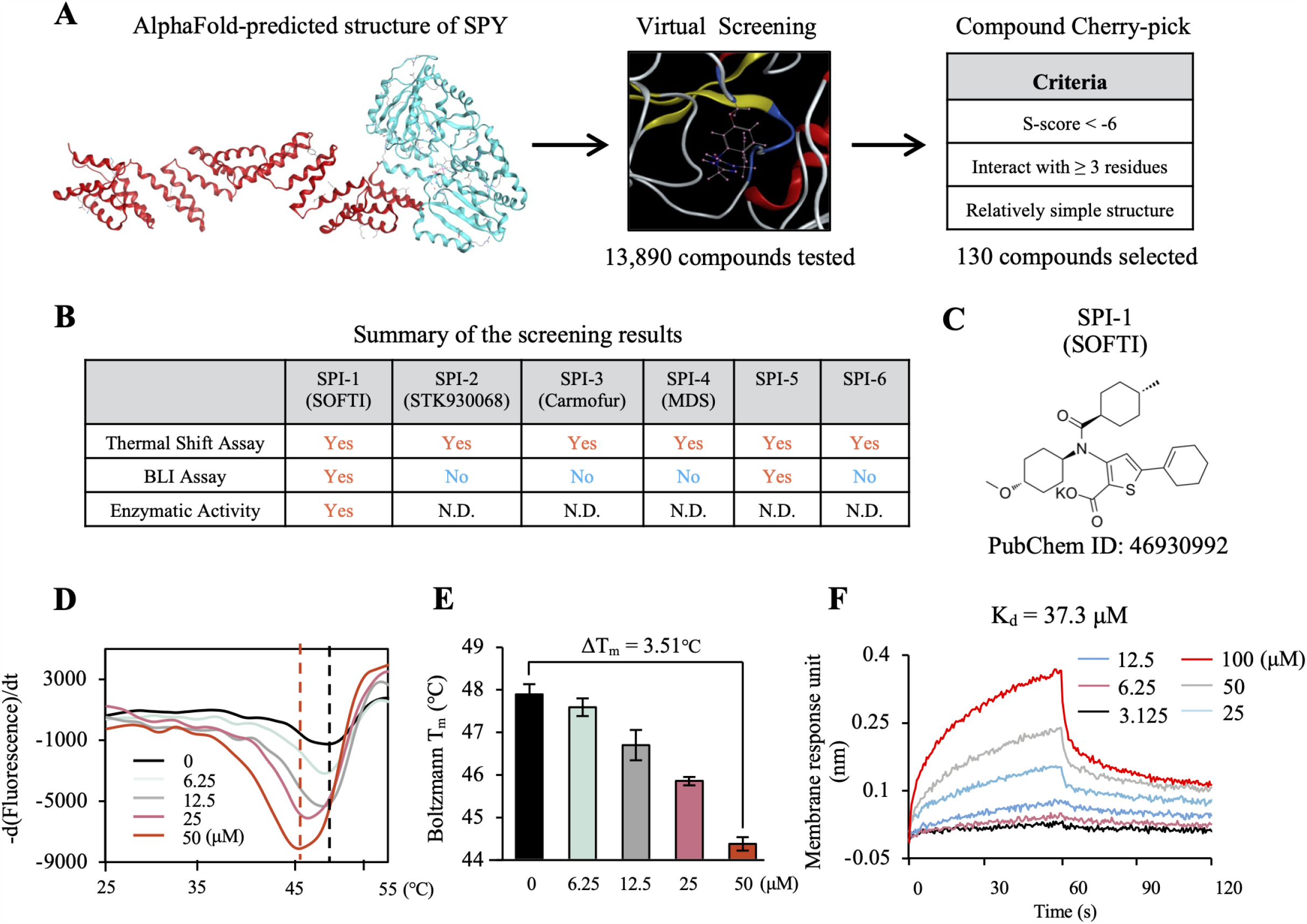
Virtual and experimental screening for SPY O-fucosyltransferase interactors. (A) A schematic illustration of primary screening with the combination of virtual docking and protein thermal shift assay to profile protein-ligand interaction. (B) Summary of the 6 compounds among 13,890 candidates that passed the three rounds of screening. (C) The chemical structure of SOFTI. (D) Protein thermal shift assay of SPY in response to varying concentrations of SOFTI (0-50 μM). (E) Quantification of the changes in melting temperature (T_m_) of SPY in response to varying concentrations of SOFTI (0-50 μM). (F) Bio-layer interferometry assay of the interaction kinetics of SOFTI with SPY.

### SOFTI inhibits the O-fucosyltransferase activity of SPY *in vitro* and *in vivo*

We tested the effects of SOFTI on SPY activity *in vitro*. SPY O-fucosylates 462 proteins including itself and the *Arabidopsis* ortholog of mammalian NEURAL PRECURSOR CELL EXPRESSED, DEVELOPMENTALLY DOWN-REGULATED GENE 1 (NEDD1), which is O-fucosylated on multiple sites (Bi et al., 2023). Therefore, we selected SPY-3TPR (N-terminal truncated SPY with three tetratricopeptide repeats) and NEDD1 as representative substrates to study SOFTI inhibition of SPY activity. SOFTI displayed concentration-dependent inhibition of the self-fucosylation of recombinant SPY-3TPR proteins, with a half maximal inhibitory concentration (IC_50_) of 13.32 μM (Figure 2A, B). Similarly, SOFTI also blocked the *in vitro* O-fucosylation of immuno-purified NEDD1-myc proteins (Figure 2C). To test whether SOFTI is a competitive inhibitor of GDP-fucose binding to SPY, we included 100 μM SOFTI and various concentrations (10, 50, 100, 200 μM) of GDP-fucose in the enzymatic reaction. In the absence of SOFTI (DMSO only as the mock control), the effect of increasing GDP-fucose concentration on SPY O-fucosylation saturated at 100 μM, with no further increase at 200 μM of GDP-fucose (Figure 2D). However, in the presence of SOFTI, which reduced the overall O-fucosylation level, 200 μM GDP-fucose further increased the O-fucosylation level compared to 100 μM GDP-fucose (Figure 2D), consistent with SOFTI competing with GDP-fucose for the substrate-binding pocket of SPY.

**Figure 2.**
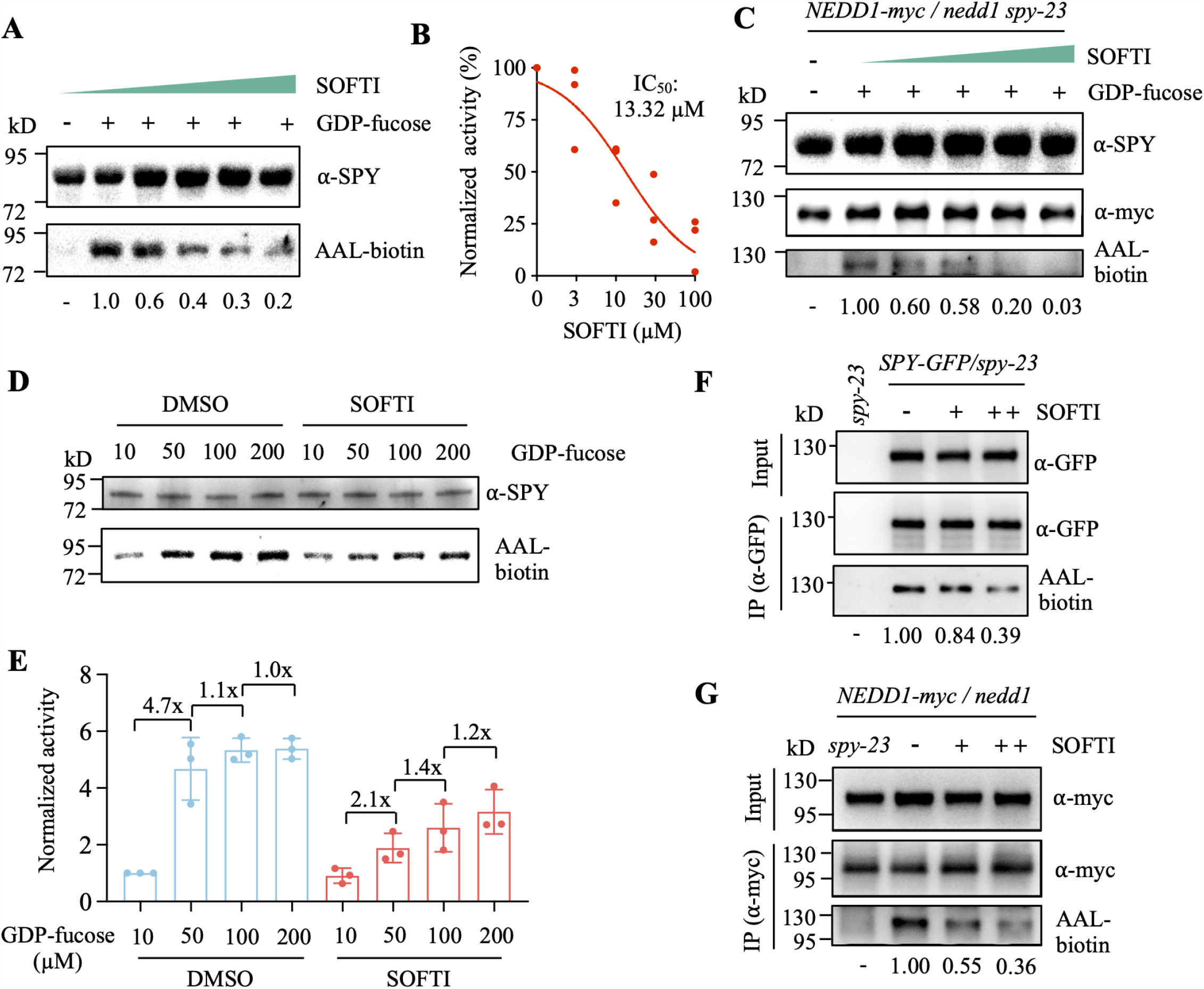
SOFT1 inhibits the enzymatic activity of SPY *in vivo* and *in vitro*. (A) Immunoblots showing the effects of SOFTI on the self-fucosylation of *E*.*coli* purified SPY-3TPR proteins with the addition of 50 μM GDP-fucose. The SOFTI concentrations are 0, 3, 10, 30, 100 μM. *The number below the lane represents band quantification, with the signal intensity of the lane without SOFTI relative to mock defined as 1*. (B) Determination of IC50 value of SOFTI via non-linear regression fitting, three points shown in the plot stands for three biological replicates. (C) Immunoblots showing the effects of SOFTI on the *in vitro* O-fucosylation level of NEDD1-myc proteins. Proteins were extracted from 12-day-old *NEDD1-myc/nedd1 spy-23* seedlings grown on 1/2 MS medium and incubated with purified SPY-3TPR proteins *in vitro* for 90 minutes with the addition of 50 μM GDP-fucose. The SOFT1 concentrations are 0, 3, 10, 30, 100 μM. (D) Immunoblots showing the competing effects of SOFTI on the self-fucosylation of *E*.*coli* purified SPY-3TPR proteins with the addition of GDP-fucose of indicated concentrations. The SOFTI concentration is fixed at 100 μM. (E) Quantification of data shown in (D), n = 3 biological replicates. (F) & (G) Immunoblots showing the effects of SOFT1 on the O-fucosylation level of SPY-GFP or NEDD1-myc proteins extracted from 12-day-old transgenic *SPY-GFP/spy-23* (F) or *NEDD1-myc/nedd1* (G) seedlings grown on 1/2 MS medium and treated with liquid 1/2 MS medium supplemented with 30 or 60 μM of SOFT1 for another 2 days. *spy-23* and *NEDD1-myc/nedd1 spy-23* were used as negative controls.

We tested the *in vivo* inhibitory effect of SOFTI. We treated *SPY-GFP/spy-23* and *NEDD1-myc/nedd1* transgenic seedlings with 10 or 30 μM of SOFTI. We immunoprecipitated the SPY-GFP and NEDD1-myc proteins and analyzed their O-fucosylation by immunoblotting. SOFTI treatment of seedlings reduced the O-fucosylation of both SPY and NEDD1 *in vivo*, indicating that SOFTI is a cell-permeable inhibitor of SPY (Figure 2E, F).

### SOFTI treatment partially mimicked *spy* mutant phenotypes

To evaluate the bioactivity and specificity of SOFTI as a SPY inhibitor, we analyzed whether SOFTI treatment causes phenotypes similar to the *spy* mutant. First, the O-fucosylation of DELLA proteins is known to inhibit GA signaling (Zentella et al., 2017), and *spy* mutants have been reported to display early-germination phenotypes (Silverstone et al., 2007). At 24 hours after imbibition (HAI), nearly all the *spy-4* seeds germinated, whereas only about 10% of the wild-type (Col-0) seeds germinated. Adding SOFTI to the medium increased the germination rate of wild type to about 50% but had no obvious effect on *spy-4*. After 60 hours of imbibition, all the seeds germinated (Figure 3A). Another phenotype of *spy* is the increased root hair density. Wildtype seedlings treated with SOFTI exhibited higher root hair density than untreated seedlings, and this response was not displayed by the *spy-4* mutant (Figure 3B, C). We recently reported that the *spy* mutant is defective in sugar-promoted seedling growth in the dark (Xu et al., 2021; Bi et al., 2023). Sugar (1% sucrose) increased the growth of wild-type seedlings after transferring from light into darkness. The *spy* mutant seedlings showed an impaired growth response to sugar. When treated with SOFTI, the wildtype seedling showed a reduced response to sugar, similar to *spy* in the absence of SOFTI. SOFTI did not affect the sugar growth response phenotype of *spy* (Figure 3D, E). Together, these results show that treatment of wildtype *Arabidopsis* with SOFTI causes *spy-* like phenotypes, whereas the *spy-4* mutant is insensitive to SOFTI treatment. These results indicate that SOFTI inhibits SPY specifically *in vivo*, and has no obvious SPY-independent, off-target effect on plant growth.

**Figure 3.**
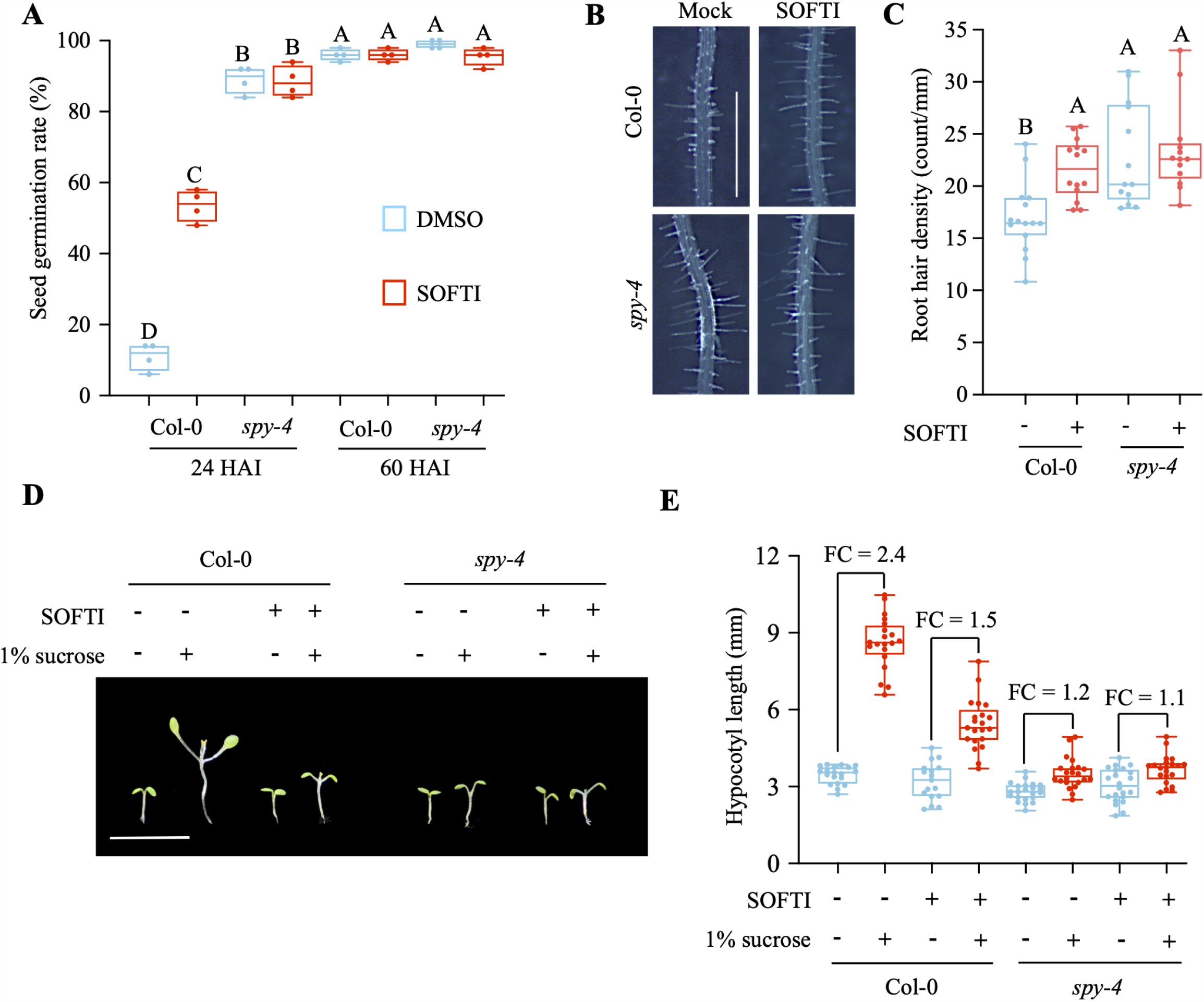
SOFT1 treatment partially mimicked *spy* mutant phenotype. (A) Seed germination rates of Col-0 and *spy-4* mutant at 24 HAI and 60 HAI with or without 30 μMSOFTI treatment. Boxes with different letters represent responses that are significantly different at P < 0.01, n = 4 for four biological replicates each containing 100 seeds. HAI stands for hours after imbibition. (B) root hair phenotype of 8-day-old Col-0 or *spy-4* seedling grown on ½ MS (1% mannitol) medium supplied with or without 10 μMof SOFTI. Bar = 1 mm. (C) Statistical analysis of root hair density (number of root hair per millimeter root length) of seedlings shown in (B), Box with different letter is significantly different at P < 0.01. (D) Phenotype of Col-0 and *spy-4* seedlings grown in long-day condition for 4 days then moved to darkness for 4 days, in the presence or absence of 1% sucrose and 30 μM of SOFTI. 1% mannitol and DMSO are used for mock control. Bar = 1 cm. (E) Statistical analysis of the phenotype shown in (D), n ≥ 15 seedlings.

### SOFTI interacts with conserved residues of the SPY substrate-binding pocket

We analyzed how SOFTI interacts with specific residues within the solved structure of the substrate-binding pocket of SPY. For our screen, we used a structure predicted by AlphaFold; however, the structure of SPY has recently been determined by crystallography and cryo-EM (Zhu et al., 2022; Kumar et al., 2023). We performed molecular docking between SOFTI and the crystal structure of SPY. SOFTI was docked to the substrate-binding pocket of SPY within the C-terminal catalytic domain (Figure 4A). A site view of molecular-docked SOFTI into SPY showed that the electronegative atoms of SOFTI form several hydrogen bonds with amino acid residues in the substrate-binding pocket. Specifically, the annular sulfate atom acts as a hydrogen bond donor to form a hydrogen bond with N661 at 3.89 Å; the carbonyl oxygen, and the two carbonyl oxygen atoms acts as hydrogen bond acceptors to form hydrogen bonds with S496, T747, K665, and G746 at 2,87, 3.29, 2.81, and 3.36 Å, respectively; and the sulfate-containing, five-membered ring forms aromatic π-hydrogen interactions with N662 at 4.53 Å (Figure 4B, C). Superposition of SOFTI and GDP-fucose showed that they partially overlapped within the pocket (Supplementary Figure 2A). SOFTI, a structurally simpler molecule, overlapped with the phosphate-phosphate-fucose moiety of the donor substrate GDP-fucose, but not the guanosine part. The three amino acid residues N662, K665, and T747 involved in SOFTI-SPY interaction also form hydrogen bonds with the pyrophosphate group of GDP-fucose according to the cryo-EM structure of SPY-GDP-fucose complex (Kumar et al., 2023). In the *spy-19* allele, K665 is mutated. We expressed SPY-K665A mutant protein in E. coli and used it *in vitro*. SPY-K665A showed no detectable O-fucosyltransferase activity (Figure 4D) and barely detectable binding to SOFTI (K_d_ = 1070 μM, compared to 41.7 μM for wild-type SPY) (Figure 4E, F). The results indicate that SOFTI mimics GDP-fucose and competes for the substrate-binding pocket of SPY, as suggested by the results of our competition assays (Figure 2D).

**Figure 4.**
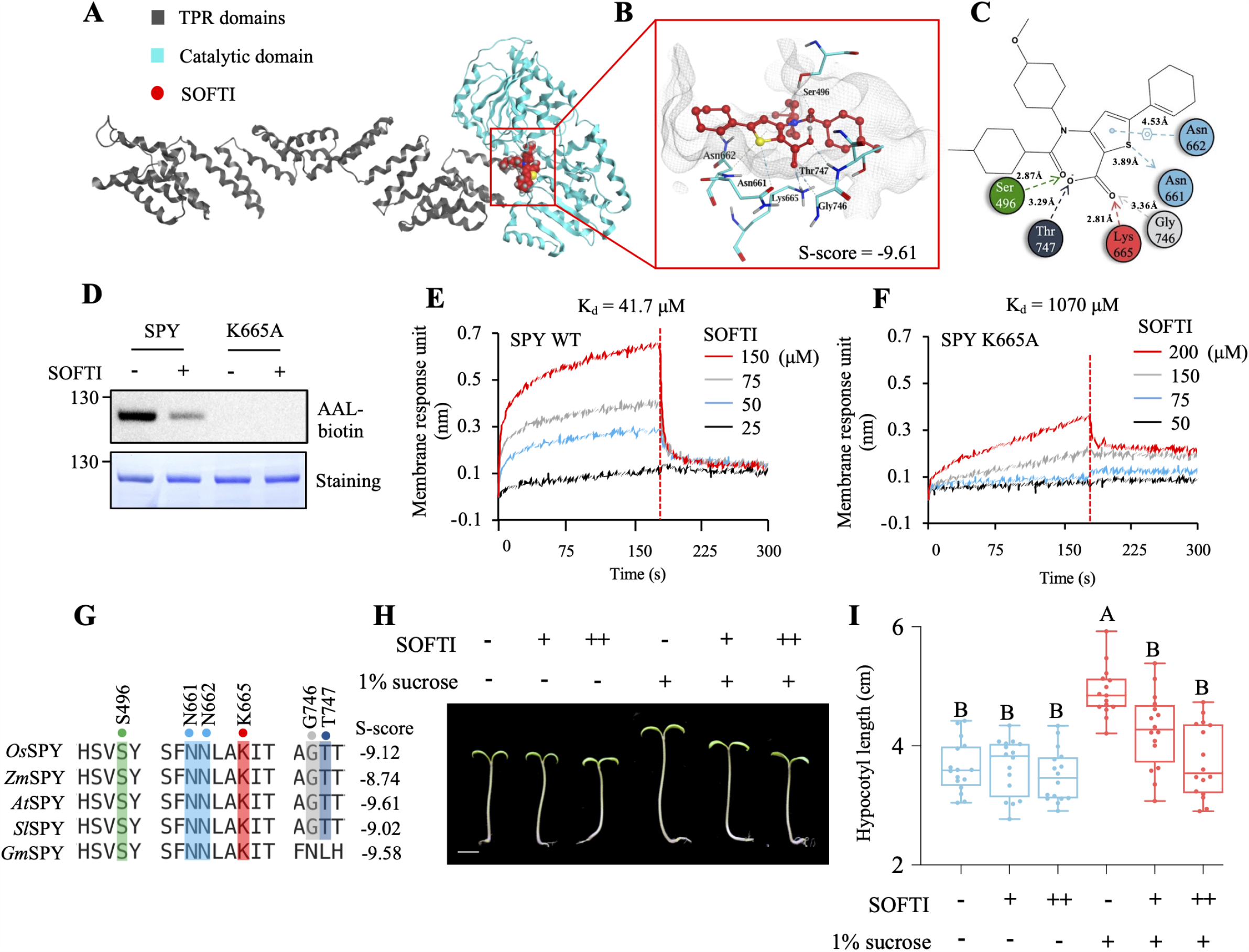
Docking simulation of SOFTI-SPY interaction. (A) A site-view of SPY and SOFT1 based on a docking simulation. (B) Detailed structure of the binding pocket of SPY (key residues shown in cyan) occupied by SOFT1 (in red), gray lines sketch the ligand binding pocket of SPY. (C) A two-dimensional illustration of the interaction between SOFTI and the amino acid residues in the SPY ligand-binding pocket in (B). (D) Immunoblots showing the enzymatic activity of SPY-WT and SPY-K665A mutant with or without 300 μM SOFTI treatment. (E) Bio-layer interferometry assay on the interaction kinetics of SOFTI with SPY-WT. (F) Bio-layer interferometry assay on the interaction kinetics of SOFTI with SPY-K665A mutant. (G) Multiple sequence alignment displaying conserved amino acid residues within the ligand binding pocket of SPY in *Arabidopsis thaliana* and its orthologs in *Oryza sativa* (*Os*), *Zea mays* (*Zm*), *Solanum lycopersicum* (*Sl*), and *Glycine max* (*Gm*). The alignment is performed with Clustal Omega. The numbers on the right panel indicate the predicted S-score between SPY orthologs and SOFTI; the lower the score, the higher predicted binding affinity. (H) Sugar-dependent growth response of tomato (*Solanum lycopersicum*) seedlings grown on 1/2MS medium with or without the addition of 1% sucrose and 10 μMSOFTI. The seedlings were grown under light for 3 days followed by 2 days in the dark. 1% mannitol and DMSO are used for mock control, respectively. Bar = 1 cm. (I) Statistical analysis of the hypocotyl lengths showed in (H), n = 16 seedlings, box with different letter is significantly different at P < 0.01.

SPY belongs to a highly conserved GT41 family of glycotransferases in higher plants (Olszewski et al., 2010). Multiple sequence alignment of *Arabidopsis* SPY orthologs in several crop species, including *Oryza sativa* (*Os*), *Zea mays* (*Zm*), *Solanum lycopersicum* (*Sl*), and *Glycine max* (*Gm*) showed that the key amino acid residues involved in SPY-SOFTI interaction are highly conserved across different species (Figure 4G). Docking simulation between SOFTI and the AlphaFold-predicted structures of SPY orthologs from crops showed comparable binding affinity to *At*SPY (Figure 4G). Consistent with the structural prediction, SOFTI inhibited the sugar-dependent growth of tomato (*Solanum lycopersicum*) seedlings in darkness but had no significant effects on seedlings grown on media without sugar (Figure 4H, I), similar to the observation in Arabidopsis (Figure 3D). These results suggest that the catalytic pocket of SPY is highly conserved and that SOFTI can inhibit SPY in different plant species.

### SOFTI derivatives screen identifies SOFTI-D1 and SOFTI-D20 as alternative SPY inhibitors

Docking simulation showed that SOFTI occupies the part in the SPY substrate-binding pocket that originally binds to the phosphate-phosphate-fucose moiety of GDP-fucose, but not the spaces occupied by the guanosine (Figure 4D). Therefore, we searched for commercially available compounds that possess the structures of SOFTI that interact with SPY and additional bulkier structures that better occupy the substrate-binding pocket of SPY. We found 21 derivatives of SOFTI and named them SOFTI-D1 to SOFTI-D21 (Supplementary Figure 3). The derivatives were docked into SPY’s substrate-binding pocket and sorted based on their affinity score (the lower the score, the stronger the predicted interaction). Four of these derivatives were predicted to have higher SPY-binding affinity than SOFTI (Figure 5A). We also tested the effects of all 21 derivatives on the self-fucosylation of SPY-3TPR, and found two, SOFTI-D1 and SOFTI-D20, that showed stronger inhibition of SPY self-fucosylation than SOFTI (Figure 5B, C). SOFTI-D1 and SOFTI-D20 also showed stronger binding to SPY in BLI assays, with K_d_ = 17.4 μM and 2.1 μM respectively (Figure 5D, E). We, therefore, performed molecular docking and superposed SOFTI-D1/D20 with GDP-fucose to understand the structural basis for inhibitor binding. SOFTI-D1 appears to occupy the same site in the pocket as SOFTI, but SOFTI-D20 competed with the GDP part of the donor substrate GDP-fucose (Supplementary Figure 2B, C). Although both derivatives showed more prominent inhibitory effects *in vitro* than SOFTI (Figure 5F), their effects on the *in vivo* O-fucosylation of NEDD1 appears comparable to SOFTI (Figure 5G). This is possibly due to their lower cell permeability compared to SOFTI. According to Crippen’s fragmentation (Ghose and Crippen, 1987), the Log*P* value of SOFTI-D1/D20 (3.05 and 4.09, respectively) are greater than that of SOFTI (2.58), indicating that they have lower cell permeability compared to SOFTI. Accordingly, each compound displayed a similar inhibitory effect on sugar-dependent growth compared to SOFTI (Figure 5H, I). In summary, we have thus identified two more compounds, derivatives of SOFTI, that show stronger inhibition of SPY activity *in vitro* and similar *in vivo* effects compared to SOFTI.

**Figure 5.**
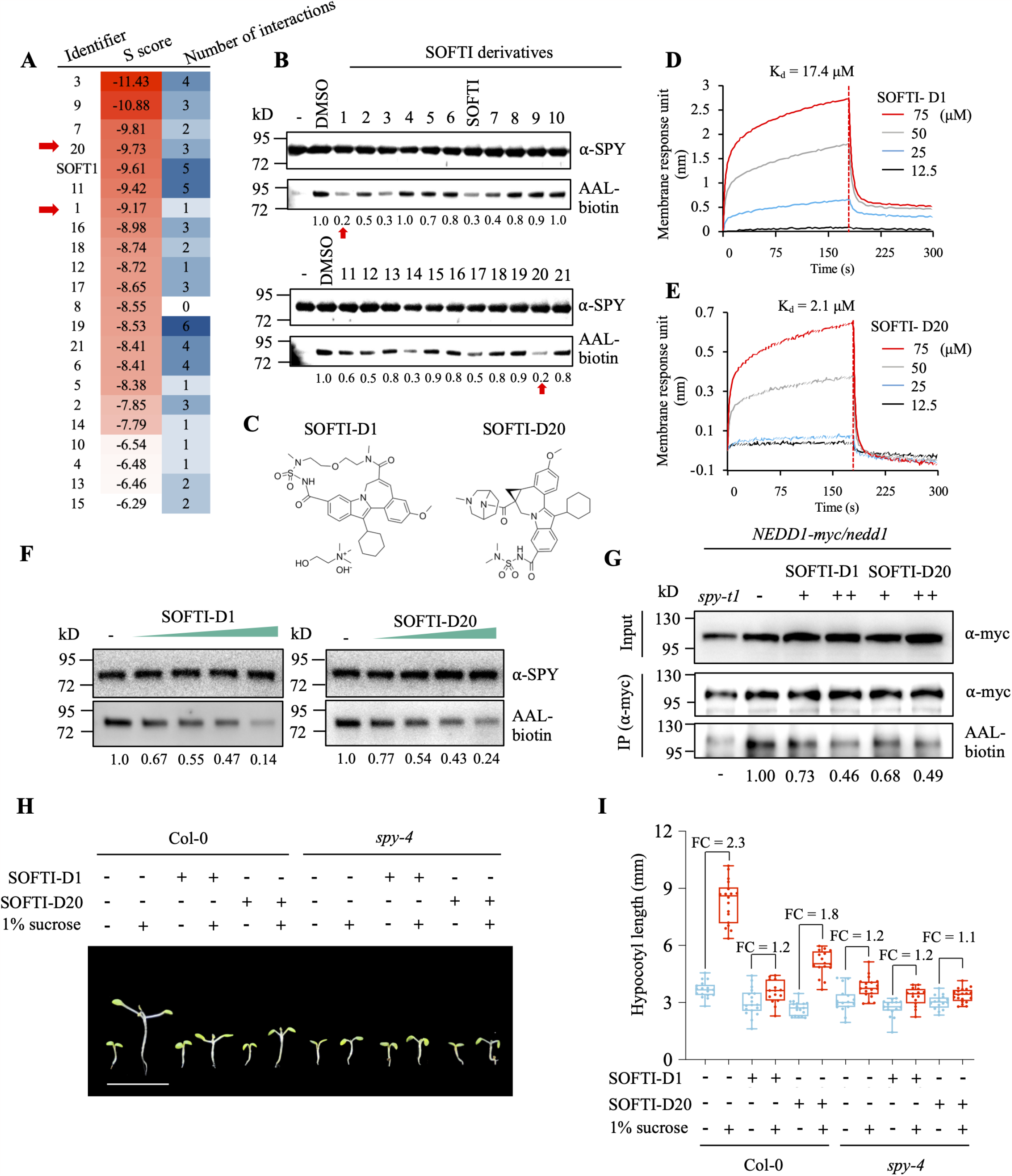
Assessment of SOFTI derivatives as SPY inhibitors. (A) Docking simulation of the interactions between SOFTI derivatives and SPY, S score indicates the theoretical affinity, lower S score indicates stronger predicted interactions. Number of interactions stands for the number of interacting amino acid residues within the ligand binding pocket of SPY. (B) Immunoblots showing the effects of SOFT1 and its derivatives on the self-fucosylation of *E*.*coli* purified SPY-3TPR proteins. The compound concentrations are fixed at 100 μM. (C) Chemical structures for SOFTI-D1 and D20, marked by red arrows in (A) and (B). (D) Bio-layer interferometry assay on the interaction kinetics of SOFTI-D1 with SPY. (F) Bio-layer interferometry assay on the interaction kinetics of SOFTI-D20 with SPY. (F) the concentration-dependent effect of SOFTI-D1&20 on the enzymatic activity of SPY-3TPR protein. The compound concentrations are 3, 10, 30, 100 μM. (G) Immunoblots showing the effects of SOFT1-D1 or D21 on the O-fucosylation level of NEDD1-myc proteins extracted from 7-day-old transgenic *NEDD1-myc/nedd1* seedlings grown on 1/2 MS medium supplemented with 10 or 30 μM of SOFT1 derivatives. *NEDD1-myc/nedd1 spy-t1* was used as a negative control. (H) phenotype of Col-0 and *spy-4* seedlings grown under long day condition for 4 days on 1/2 MS plates with 1% sucrose and transferred into liquid 1/2MS medium supplemented with 1% sucrose or 30 μM of SOFTI-D1/D20 in darkness for 5 days. 1% mannitol and DMSO are used for mock control, respectively. Bar = 1 cm. (I) Statistical analysis of the phenotype shown in H, n ≥ 15 seedlings.

## Discussion

Being an important regulator of cellular signal transduction, O-glycosylation of proteins are known to be essential for cellular homeostasis and growth regulation in both animals and plants (Bi *et al*., 2023; Hart *et al*., 2007; Zentella *et al*., 2023). While the single mutants of the *Arabidopsis* O-glycotransferases SEC and SPY show various mild developmental defects, the double mutant is embryonic lethal (Hartweck *et al*., 2002). Conditional inhibition is required to further study the function of O-glycosylation, particularly in post-embryonic development stages. Small molecule inhibitors of human OGT have been well-characterized and widely used (Martin *et al*., 2018; Ortiz-Meoz *et al*., 2015). We found, however, that these OGT inhibitors do not inhibit SPY even at high concentrations (Supplementary Figure 4), reflecting the need to develop specific inhibitors for the O-glycotransferases in plants. Therefore, using virtual screening followed by *in vitro* and *in vivo* tests, we identified SOFTI, the first small molecule inhibitor of SPY.

Virtual screening identified candidate chemicals that bind to the catalytic pocket of SPY. *In vitro* experiments, including TSA and BLI assays, demonstrated that SOFTI directly interacts with SPY (Figure 1). Further, molecular docking of SOFTI to the catalytic pocket of SPY showed that SOFTI interacts with multiple key amino acid residues that interact with the donor substrate GDP-fucose according to the crystal and cryo-EM structures (Kumar et al., 2023; Zhu *et al*., 2022) (Figure 4A-D). These include K665, which is essential for both enzyme activity and SOFTI binding (Figure 4E-F).

The SPY-inhibitor activities of SOFTI, SOFTI-D1 and SOFTI-D20 were confirmed by *in vitro* and *in vivo* assays. We show that SOFTIs inhibit SPY’s self-fucosylation activity and SPY O-fucosylation of NEDD1 in a dose-dependent manner (Figures 2 and 5). SOFTI treatments of wild-type Arabidopsis seedlings resulted in reduced protein O-fucosylation and *spy*-like phenotypes including early seed germination, increased root hair density, and sugar-insensitive growth arrest in the dark (Figures 3 and 5). Importantly, SOFTIs caused no significant phenotypic change in the *spy* mutant (Figures 3 and 5), indicating that SOFTIs are specific SPY inhibitors with little or no SPY-independent effects on plant growth.

Structure-based virtual screening greatly simplifies the candidate selection process compared to the traditional phenotype-based or assay-based high-throughput screen approaches. However, virtual screening relies on high quality three-dimensional structure of the target protein. As crystal structures remain unavailable for most plant proteins, virtual screening has not been widely used for plant research. Recently, structural predictions have become available from AlphaFold. Whether the structures predicted by AlphaFold are accurate and reliable enough for virtual screening remains unclear, as conflicting conclusions have been reported (Wong et al., 2022; Scardino et al., 2023). Our study represents a successful case, perhaps the first, of using AlphaFold-predicted protein structure to identify small-molecule inhibitors of a plant protein.

The crystal and cryo-EM structures of SPY have been recently determined (Zhu et al., 2022;Kumar et al., 2023), providing a timely opportunity to compare predicted and solved structures. Structure-based superposition of the crystal structure and the AlphaFold-predicted structure of the full-length SPY showed an overall root means square deviation (RMSD) of 9.339 Å, indicating a significant deviation between the two structures (Supplementary Figure 5A, B). However, the superposition of only the catalytic domain of SPY resulted in an RMSD of 0.665 Å between the AlphaFold-predicted and the solved crystal structures or 0.61 Å between the AlphaFold-predicted and Cryo-EM structures (Supplementary Figure 5C), which are similar to the median backbone accuracy of overall AlphaFold structures (0.96 Å RMSD_95_. Jumper et al., 2021). Consistent with the accuracy of the predicted structure and docking simulation, some residues of SPY that interact with SOFTI in the simulation were found to form hydrogen bonds with GDP-fucose in the cryo-EM structure (Figure 4D) (Kumar et al., 2023). By screening derivatives of SOFTI virtually and then experimentally, we identified SOFTI-D1 and D20 as more potent inhibitors of SPY (Figure 5). Our study illustrates how the development in computational structural biology can advance the discovery of chemical tools for plant biology and agriculture. While no crystal or Cryo-EM structure is available for most plant proteins, our results indicate that the structures predicted by AlphaFold provide a viable option for virtual screening for chemical binders. As AlphaFold predicts the structure of all proteins, virtual screening, as illustrated in our study, will have a broad impact on plant science.

SOFTI is a useful tool for functional studies of O-fucosylation in plants. For example, when combined with genetic mutants, chemical inhibitors applied spatiotemporally can uncover the post-embryonic functions shared by SPY and SEC. The rapid inhibition of SPY by a small molecule inhibitor will also enable analysis of the dynamics of molecular and cellular responses to the alteration of protein O-fucosylation *in vivo*. Furthermore, the residues in the Arabidopsis SPY ligand-binding pocket involved in GDP-fucose-binding, as well as SOFTI-docking, are conserved among higher plants. Consistent with the idea that this conservation is functionally significant, SOFTI showed a similar growth-inhibition effect in tomato as in Arabidopsis (Figure 4). SOFTI thus can be used as a selective inhibitor of O-fucosyltransferases in agriculturally important plants, where *spy* mutants are mostly lacking, enabling broad functional studies of O-fucosylation as an essential sugar-sensing mechanism. Given the important role of O-fucosylation in plant development, it is conceivable that SOFTI and its derivatives can be developed into agrochemicals that improve food production, in addition to their application in basic research of plant biology.

## Materials and Methods

### Plant materials and culture conditions

All *Arabidopsis thaliana* (Arabidopsis) materials used in this study are in Col-0 ecotype background and grown in a long-day growth chamber with 16h light and 8h dark cycle at 22°C. The seeds were surface-sterilized by immersing in 75% ethanol for 10 mins followed by rinsing with autoclaved water for three times and then placed on 1/2 Murashige and Skoog (MS) culture medium containing 1.2% (w/v) agar and supplied with 1% mannitol or 1% sucrose as indicated. The T-DNA insertion line *spy-4* (Bi et al. 2023), *spy-23* (Bi et al., 2021) and transgenic line *NEDD1-myc/nedd1* (Zeng et al., 2009) have been described previously. The *NEDD1-myc/nedd1 spy-23* was generated via genetic crossing.

### Plasmid construction

The coding sequence of full-length SPY and 3TPR-truncated SPY were amplified with primers from Table S1. The PCR amplified sequences were cloned onto the pET28sumo expression vector with a 6xHis-SUMO tag at the N-terminus. The point mutant K665A SPY was generated via Q5 Site-Directed Mutagenesis Kit (NEB, E0552S). All plasmids were sequenced to ensure accuracy.

### Protein expression and purification

The full-length or 3TPR-truncated cDNA sequence of SPY was amplified from *Arabidopsis* cDNA with appropriate primers (listed in Supplemental Table 1) and cloned into the pET28sumo vector (Addgene?), which adds a 6xHis-SUMO tag to the N-terminus of the protein of interest. The plasmid carrying SPY full length sequence or 3-TPR truncated sequence was transformed into Rosetta DE3 *E. coli* strains for protein expression. The transformed bacteria were then cultured to OD= 0.8 at 37°C and then cooled to 16°C, followed by addition of 300 μM (final concentration) isopropyl-β-D-thiogalactoside (IPTG). After overnight culture, the cell pellets were then collected and resuspended in 1xPBS (2.7 mM KCl, 2 mM KH_2_PO_4_, 137 mM NaCl, 10 mM Na_2_HPO_4_, pH = 7.4). The proteins were then affinity purified with Ni^2+^ columns (Invitrogen, R901-01), and the eluted proteins were further purified via gel filtration chromatography with a superdex 200 (GE Healthcare) column. The final protein product was stored in 1xPBS at 1 mg/mL at -80°C.

### Protein thermal shift assay (TSA)

Purified proteins were thawed on ice and mixed with 1x SPYRO Orange dye and compounds at the indicated concentration (1% DMSO final), followed by incubation on ice for 30 minutes and then put into a QuantStudio Q6 (Thermo Fisher Scientific) for the detection of fluorescence. The program gradually increases the sample temperature from 25°C to 99°C with a speed of 0.05°C/s. The derivative of the fluorescence curve was then generated with Protein Thermal Shift ™ software and Boltzmann fit was utilized to determine the T_m_ temperature of each sample. Four replicates were included for each concentration of the compounds.

### Biolayer interferometry (BLI)

The biolayer interferometry was performed as previously described (Xie et al., 2022) using Octet^®^ R2 Protein Analysis System (Sartorius, Figure 1) or Gator-plus (Gator Bio. Figures 4 and 5) with some modifications. Briefly, 100 μL of 1 mg/mL purified SPY full-length protein was biotinylated at a 1:2 (protein:biotin) ratio, and free biotin was separated with a desalting column (#G-MM-ITG, www.bomeida.com). Super Streptavidin sensors (Fortebio) were dipped into the protein solutions for 15 mins for loading, and then dipped into various concentrations of SOFTI (0-100 μM). The dissociation constant K_d_ was calculated via Fortebio analysis software based on the ratio of K_off_/K_on_ values. For Figures 4E, 4F, 5D, and 5E, BLI was performed with a similar protocol but on GatorPlus Biosensor System (GatorBio) with Small Molecule Analysis Probe (GatorBio, 160011).

### Molecular docking and virtual screening

The crystal structure of SPY was obtained from Protein Data Bank (PDB) with PDB ID: 7Y4I, and the AlphaFold-predicted structure of SPY was downloaded from https://alphafold.com. Virtual screening, protein structural superposition, and molecular docking simulations were performed with the commercial software Molecular Operating Environment (MOE) (Chemical Computing Group, 2020.09) purchased from https://www.chemcomp.com. The binding affinity of small molecules and proteins are predicted based on the Generalized Born solvation model (GBVI), which is a scoring function of the free energy of the protein bound to each molecule. For SOFTI docking with SPY protein, 1000 random poses of SOFTI were generated and docked one-by-one into the substrate-binding pocket of SPY with the induced fit model, and the pose with the lowest estimated free energy was analyzed and displayed.

### Protein immunoprecipitation and immunoblotting

Arabidopsis seedlings were grown on 1/2 MS plates with or without treatment for 12 days. 100 mg tissue of the whole plant seedling was harvested and ground into powder in liquid nitrogen. 150 μL ice-cold IP buffer (50 mM Tris-HCl, pH = 7.5, 50 mM NaCl, 10% glycerol, 0.25% Triton-X 100, 0.25% NP40, 1 mM PMSF, 1x Halt™ protease inhibitor cocktail (Thermo Scientific), 1x Pierce™ phosphatase inhibitor (Thermo Scientific) was added into the powder and lysed on ice for 10 mins. The samples were then ultrasonicated for 1 min with 5s on/ 5s off cycle to release nuclear proteins and centrifuged at 14, 000 rpm for 15 mins at 4°C. 20 uL anti-myc (Thermo Scientific, 88843) or anti-GFP (SMART lifesciences, SM038005) magnetic beads were then incubated with the supernatant for 1 hour. After two rounds of washing with IP buffer, the proteins were eluted from the beads via adding 2xSDS loading buffer (60 mM Tris–HCl, pH 6.8, 25% glycerol, 2% SDS, 14.4 M β-mercaptoethanol, 0.1% bromophenol, and 1 M dithiothreitol) and boiling at 95°C for 10 mins.

For immunoblotting, the protein samples were loaded onto 10% polyacrylamide gels and then transferred onto polyvinylidene difluoride (PVDF) membrane. The membrane was blocked with 3% BSA to exclude non-specific binding, and then probed with anti-myc (Cell Signaling Technology, 9B11, 1:2000 dilution), anti-GFP (Transgene, Q20329, 1:2000 dilution), anti-SPY (PhytoAB, PHY1737S), or Biotinylated Aleuria aurantia lectin (AAL, Vector Laboratories, ZF0305) with 1:2000 dilution in 3% BSA. After three rounds of washing with 1xPBST (0.1% Tween-20), 6 mins each, the membrane was then probed with anti-mouse, anti-rabbit secondary antibodies (BIO-RAD, 1:5000 dilution) or Pierce™ HRP-conjugated Streptavidin (Thermo scientific, 1:2000 dilution).

### SPY in vitro O-fucosyltransferase assay

To determine the O-fucosyltransferase activity of SPY, 5 μg of SPY-3TPR recombinant protein (for the case of NEDD1-myc protein, 5 μg SPY-3TPR protein and 5 μL of anti-myc beads were added together) was added into a 30 μL reaction volume containing 1xPBS, 5 mM MgCl_2_ with the compounds at indicated concentration (DMSO 1% v/v). The reagents were incubated on ice for 30 mins for proper binding of the small molecules to SPY. Then 50 μM (final concentration) of GDP-fucose was added, and the tubes were placed on a rotator for 1.5 hours at 22°C. The reaction was terminated with the addition of 2xSDS loading buffer and boiling at 95°C for 10 mins.

## Supplementary data

**Supplementary Figure 1. Purification of SUMO-tagged SPY full length and SPY-3TPR proteins**. (A) Coomassie brilliant blue stained SDS-PAGE with purified SPY full length protein and SPY-3TPR truncated protein.

**Supplementary Figure 2. SOFTI-D1 and SOFTI-D20 overlap with different moieties of GDP-fucose**. (A) A site view of the ligand binding pocket (sketched by grey lines) of SPY containing molecular docked SOFTI (in red) superposed to GDP-fucose (in green), key amino acid residues in the ligand binding pocket are shown in cyan. (B) A site view of the ligand binding pocket (sketched by grey lines) of SPY containing molecular docked SOFTI-D1 (in purple) superposed to GDP-fucose (in green), key amino acid residues in the ligand binding pocket are shown in cyan. (C) A site view of the ligand binding pocket (sketched by grey lines) of SPY containing molecular docked SOFTI-D20 (in blue) superposed to GDP-fucose (in green), key amino acid residues in the ligand binding pocket are shown in cyan.

**Supplementary Figure 3. Chemical structures of SOFTI derivatives**. (A) The chemical structure and PubChem ID of the SOFTI derivatives used in this study

**Supplementary Figure 4. OGT inhibitors do not interfere SPY enzyme activity**. Detailed structure of the binding pocket of SPY (in cyan) occupied by OSMI-1 (A) and BADGP (B) based on molecular docking simulation, OSMI-1 is in orange and BADGP is in purple. (C) immunoblots showing the effects of OGT inhibitors and SOFT1 on the self-fucosylation of *E*.*coli* purified SPY-3TPR proteins with the addition of 50 μM GDP-fucose. “+” and “++” represents 100 and 300 μM respectively.

**Supplementary Figure 5. Structural superposition of AlphaFold-predicted SPY structure and SPY crystal structure**. (A) a schematic diagram showing the domains of SPY protein. (B) structural superposition of the SPY crystal structure (PDB code 7Y4I, in cyan) and AlphaFold-predicted structure of SPY (uniport ID Q96301, in red) based on the full-length sequence, RMSD stands for root mean square deviation. (C) structural superposition of the SPY crystal structure and AlphaFold-predicted structure of SPY based on the structure of the catalytic domain (amino acids 431-825)

## Acknowledgements

We thank Dr. Bo Liu from University of California, Davis for sharing the NEDD1-myc/nedd1 transgenic lines. This work is supported by a grant from the National Institute of Health to Z-Y.W (R01GM066258) and grants from National Natural Science Foundation of China (Grant No. 21907049 to K.J and Grant No. 3191154007091740203 to H.G).

## Author Contributions

Z.-Y.W. and Y.A. designed the research; Y.A., H.Z., and K.J. performed the virtual chemical screening and docking simulations; Y.A., Q.Y. and G.G. performed the experimental chemical screening and ligand-protein binding assay with supervision by H.G. and K.J.; Y.A. Q.Y., and G.G. purified the proteins for enzymatic assays; Y.A. performed the enzymatic assays with assistance from Y.B., Z.Z., and H.Z.; Y.B. and Z.Z. constructed the transgenic plant materials; Z.-Y.W. supervised the project; Z.-Y.W. and Y.A. wrote the manuscript with input from all co-authors.

## Data Availability

All data presented in this study are available from the corresponding author on request.

